# Automated Seizure Detection Tailored Towards Portable Patient Care Using A Novel, Computationally Efficient Method of Analysis

**DOI:** 10.1101/495572

**Authors:** Zachary Huang, Yiming Ying

## Abstract

Epilepsy is a devastating neurological disorder that affects approximately 1-3% population worldwide. Wearables have recently gained more popularity as having a promising future in epilepsy management including seizure alert and close-loop therapy for severe forms of seizures. Due to the random and low frequency of seizures, seizure evaluation requires continuous, long-term EEG monitoring, which produces a large volume of data. The future treatment systems will rely on algorithms that can detect seizures with high precision and low computational cost. Previous implemented algorithms have used various computationally expensive methods of data transformation and feature extraction. In the present study, we developed a new computationally efficient seizure detection algorithm based on analysis of the broad, global shape EEG data of tonic-clonic seizures from a mouse model of temporal lobe epilepsy (TLE). To perform this algorithm, EEG data was normalized and processed through a rolling mean function, producing smoothed, simplified EEG clips that represent the global shape of the clip. These signals were then directly inputted for SVM training and testing. This novel method of seizure analysis only requires a small fraction of EEG data points, yet achieved an accuracy rate of approximately 98.51%. Our study provides a proof of principle that this simpler method could have an advantage in low-power platform such as wearables. Recently, the FDA approved a seizure detection app called embrace2 by *Empatica*. However, their algorithm detects seizures indirectly by monitoring heart rate and muscle contractions. Our novel algorithm detects seizures directly through analysis of brain activity. Thus, our algorithm may be better suited for future wearables in epilepsy management.

## Introduction

Epilepsy is a devastating neurological disorder that affects approximately 65 million people worldwide (Sauro et al., 2016). Epilepsy has been linked to an increase in morbidity and mortality (Levira et al., 2017). In addition, approximately one-third of patients with epilepsy are drug-refractory, and thus still experience seizures. Moreover, status epilepticus is a severe life-threatening form of seizure event which consists of multiple seizures in succession, lasting for a prolonged period of time. This type of seizure event is especially dangerous, and requires fast detection and rapid, on-demand treatment. There is still an urgent for fast, reliable detection of seizures with reduced response time for treatment.

Electroencephalography (EEG) recording has become an essential tool in evaluating seizure activity, critical for both epilepsy research and treatment. However, due to the randomly occurring and infrequent nature of seizures, seizure evaluation requires continuous, long-term EEG monitoring, which produces a huge amount of data. Analysis of this EEG data is extremely burdensome and labor-intensive, often involving manual visual scanning of large EEG records by trained personnel. Computer-aided detection systems have been explored since the early 1970s (Baldassano et al., 2017; Saini and Dutta, 2017). Lately, many automated detection systems based on machine learning algorithms have been proposed (Feng et al., 2018; Liu et al., 2012; Saini and Dutta, 2017; Zhang and Chen, 2017). A plethora of feature extraction techniques have been implemented, from Fourier and wavelet transformations to non-linear dynamics such as fractal dimensions. These techniques, combined with machine learning classifiers such as support vector machine and artificial neural networks, have produced promising results in both seizure detection and prediction. However, these programs are still unsatisfactory for deployment in portable patient care, such as wearables, due to their high computational cost.

Closed-loop therapy seems to be a promising future in epilepsy treatment. However, it relies on fast and accurate detection or prediction of seizures on low-power devices such as wearables. While seizure prediction may seem like a more promising endeavor than seizure detection, current prediction models have been inapplicable in closed-loop therapy due to their high computational cost. The aim of this project is to develop a novel seizure detection algorithm with low computational cost and high accuracy, applicable for wearable use. Trained technicians and physicians can identify seizure based on their global shape. In contrast, the algorithms that have been developed in the previous studies have analyzed minute fragments within the data for discrete, non-visual features within the data. Inspired by the ability of trained personnel to effectively identify seizures without analyzing minute features of seizure events, this present study attempts to model the global approach of seizure detection. We optimize detection of tonic-chronic seizures in pilocarpine model of temporal lobe epilepsy (TLE), the most common form of human epilepsy and most similar structurally to status epilepticus. By using very simplistic methods of data processing to generate the global shape of each seizure, we are able to achieve over 98% accuracy and over 96% sensitivity and 98% specificity while significantly minimizing the computational cost.

## Material and Methods

### I. Materials

EEG data was provided by an epilepsy laboratory at Albany Medical College. The EEG data for spontaneous recurrent seizures was collected via epidermal recording of 13 mice that were treated by pilocarpine, which models temporal lobe epilepsy (TLE). Continuous recording of EEG data lasted for 24 hours a day, 7 days a week, for two months, amassing over 18,000 hours of EEG data. Seizures were manually identified and verified by lab technicians in corroboration with video recording. 32 seizure clips and 32 non-seizure/noise clips of ∼60 seconds long were split randomly for training and testing groups for cross-validation. In addition, 546 similar-length clips of seizures and non-seizure/noise events were used for final assessment of the algorithm. Seizure events are relatively rare in frequency in comparison to non-seizure/noise events. To attempt to model real scenarios, we set the seizure-to-non-seizure/noise ratio of our final assessment data as 1:13.

An HP Z840 Workstation with 128 GB RAM was used to harvest the EEG data clips. Data transformation, SVM algorithm training, and testing were done on a 2016 MacBook Pro with 8GB RAM in the Python development environment Spyder. The packages Pandas, Numpy, Matplotlib were used for data transformation. The package Scikit-Learn was used for its adaptation of SVM in Python. SVM was used for its effectiveness in high dimension-to-sample ratio, and for its relatively low memory cost.

### II. Data Pre-Processing

The schematic diagram in **Figure 1** shows the overall strategy of data processing, SVM algorithm training and testing. EEG data was first trimmed of extraneous labels and timestamps. Then, each EEG clip was individually processed through the standard data normalization formula:

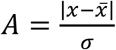

**Figure 1:**
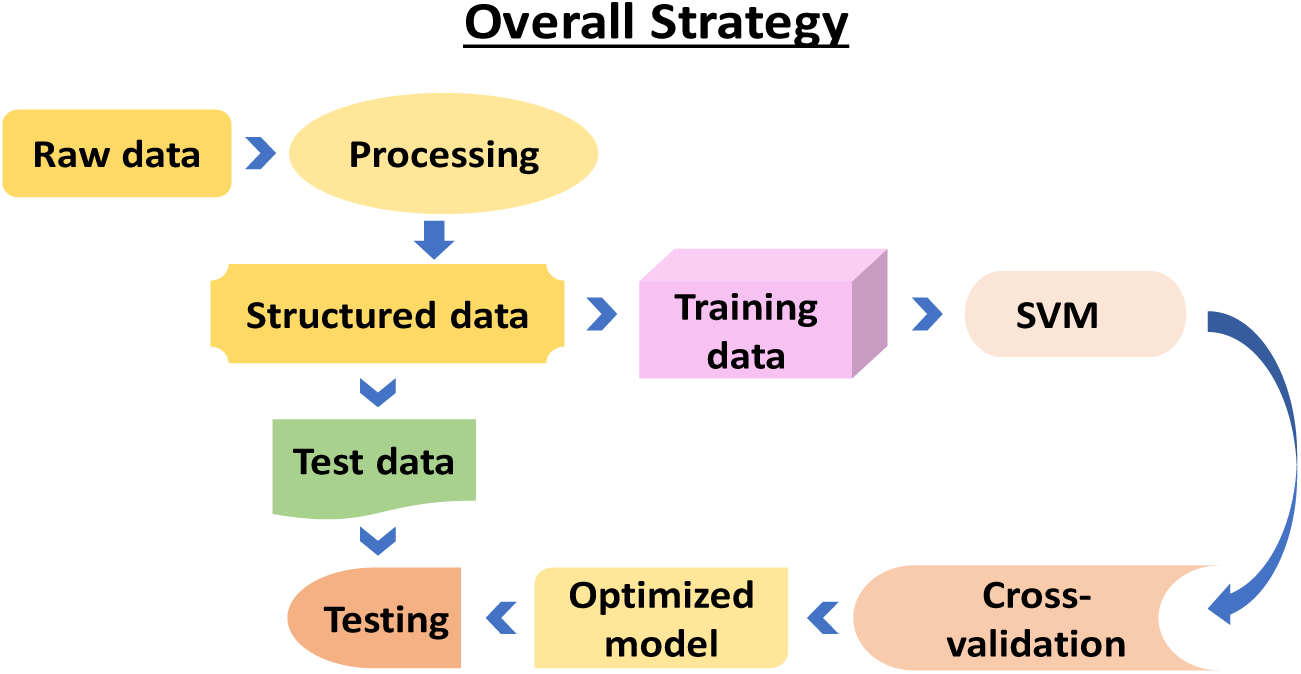
Overall strategy of EEG data processing and seizure classification.

For each clip, the mean and standard deviation of the individual clip was taken. Then each value in the clip was subtracted by the mean and divided by the standard deviation. Because seizure clips inherently have higher standard deviations, the same baseline value in both the seizure and non-seizure clip would be decreased in the seizure clip, as it would be divided by a higher number. After normalization, the dataset was inputted into a simple moving average formula, defined as:

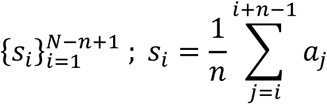

This data transformation smoothed the data in each clip, allowing for seizure and non-seizure clips to be simplified into their general shape. Because our method looks at the global shapes, there is forgiveness in the specificity and detail of the data. Not all points in each clip are necessary for robust analysis, as analysis of the general shape does not require such specificity. Thus, after each clip was transformed, each clip was normalized in length to 1000 points to save computational cost. For a dataset consisting of *p* points, WEtook every *n*th point to cut it down to 1000 points, where *n* is defined as:

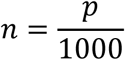

This method dramatically lowers computational cost, cutting each clip from 10,000+ points to 1000, while preserving the shape for SVM analysis. Afterwards, the entire dataset was divided by the max of the dataset to condense all values to between 0 and 1. This was necessary to feed the data into the SVM algorithm, as the SVM only works with vectors of identical length and with values between 0 and 1. As indicated in **Figure 2**, data processing results in general shape of seizure event, which is quite different from the shape of noise.

**Figure 2:**
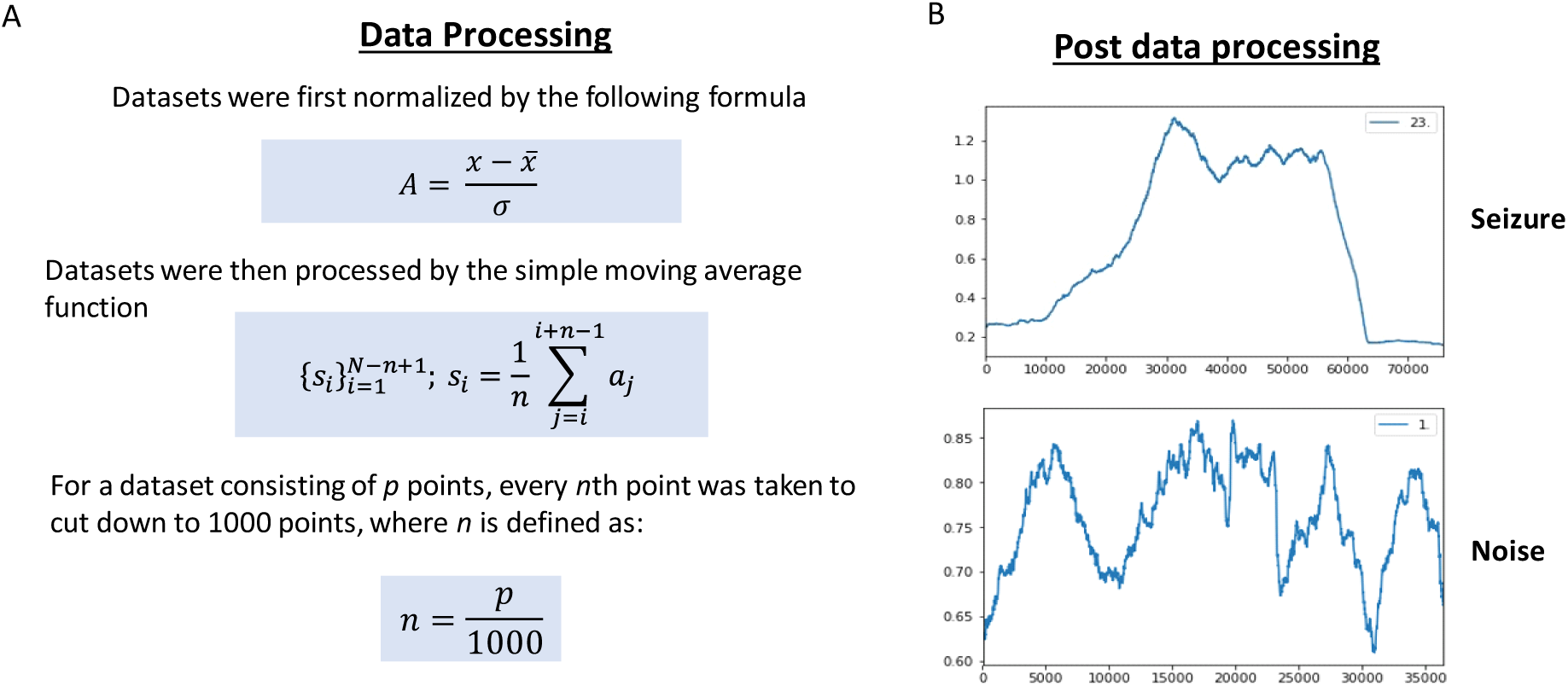
A) EEG data sets underwent multiple steps of transformation. B) Example of data after transformation. Seizure and noise signals were all normalized and smoothed out to represent general data shape.

### III. SVM Algorithm Training

The kernel-based SVM built on statistical learning theory was developed for binary classification (Cortes and Vapnik, 1995). The idea of the SVM algorithm is to project nonlinear separable samples onto higher-dimension spaces using kernel functions and then create the optimal separating hyperplane in the higher-dimension space (**Figure 3**). The optimal separating hyperplane is computed by solving a quadratic optimization problem. In this experiment, the type of kernel function used was a radial basis function (RBF), defined by the following equation:

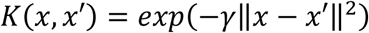

**Figure 3:**
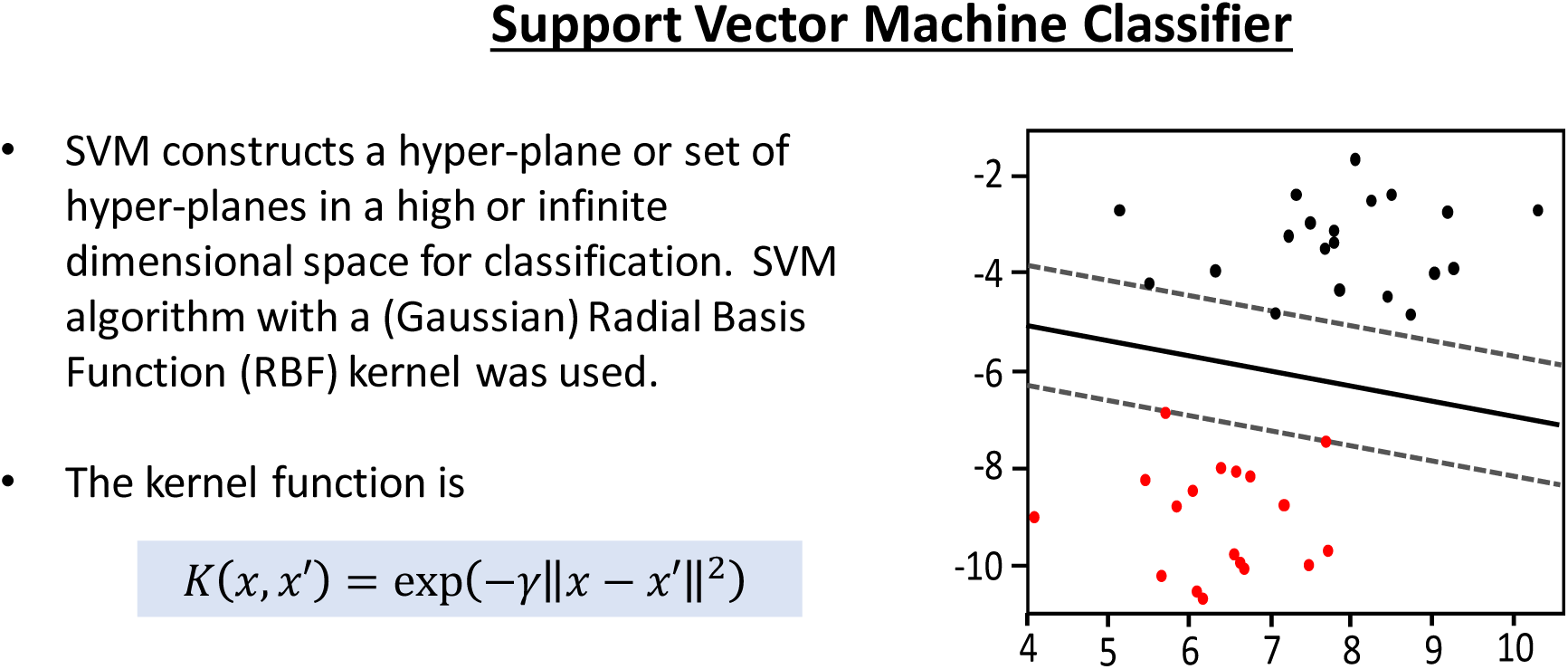
The diagram illustrates the statistical learning theory of the kernel-based SVM.

However, the RBF kernel in SVM requires 2 parameters, C and gamma. Intuitively, the gamma parameter defines how far the influence of a single training sample reaches. The C parameter trades off misclassification of training examples against simplicity of the decision surface. A low C makes the decision surface smooth, while a high C aims at classifying all training examples correctly by giving the model freedom to select more samples as support vectors.

The optimal parameters for the best model can be found by performing the standard cross-validation. One round of cross-validation involves partitioning data into complementary subsets, performing the analysis on one subset (called the training set), and validating the analysis on the other subset (called the validation set). To reduce variability, in most methods multiple rounds of cross-validation are performed using different partitions. Specifically, for the 10-fold cross validation, the training data was randomly split into 10 parts, and for each value of C in the set (0.001,1000) and each value of gamma in the set (0.001,1000), 9 parts of the data are trained, and each combination of parameters are tested with the last portion of the split dataset. The error on each test is recorded, and the C and gamma values that correspond to the highest accuracy rate are kept. This allows for an exhaustive search for the best parameters (**Figure 4**).

**Figure 4:**
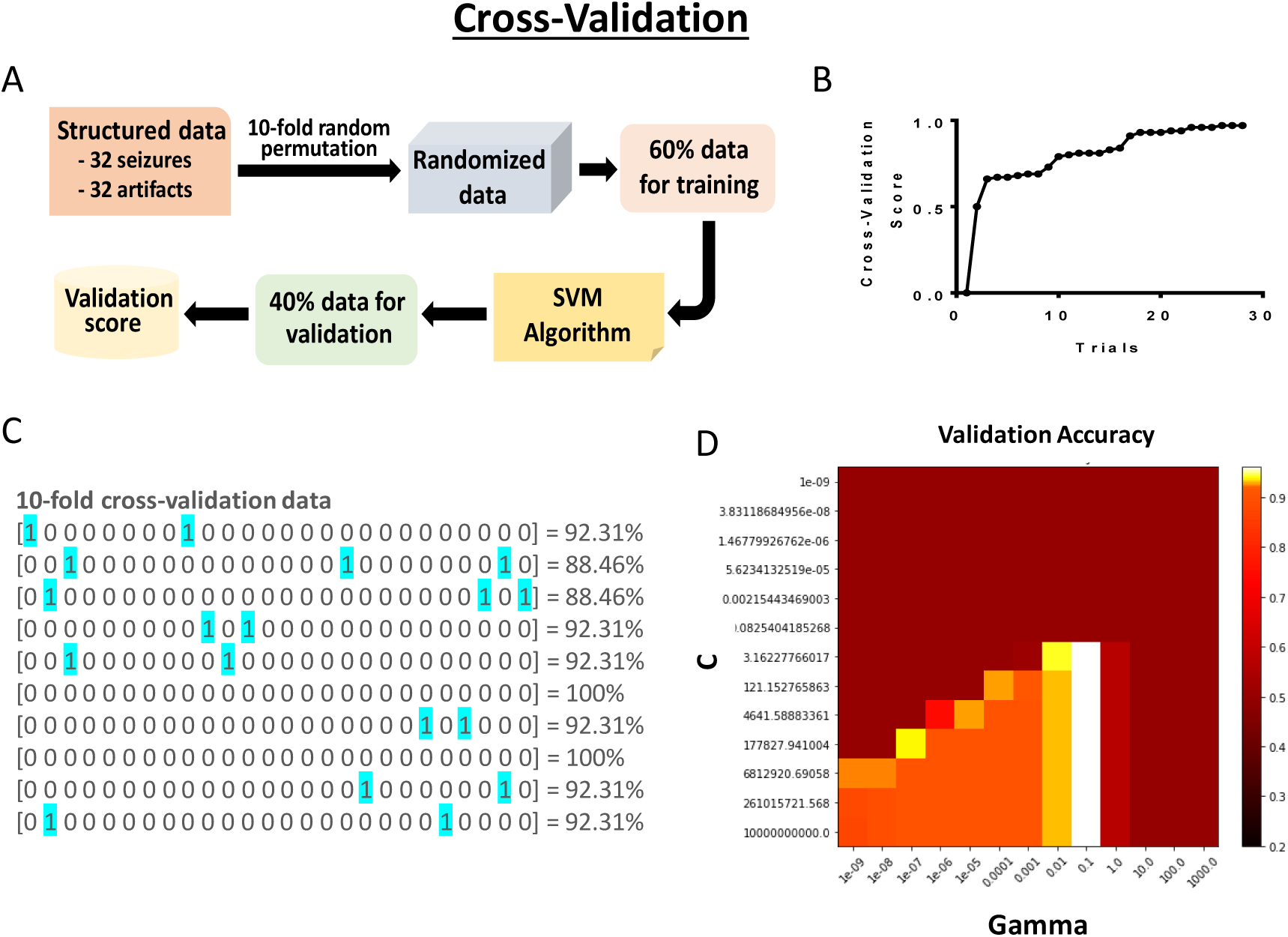
A) Schematic diagram showing the steps of cross validation. B) Improvement of cross-validation score over the course of trials. C & D) Cross-validation results. The best parameters are (‘C’: 3.162, ‘gamma’: 0.100) with a score of 0.96.

After the optimal parameters were found, the algorithm model was tested on 546 EEG clips which contained 42 seizures events and 504 non-seizure events. Accuracy was recorded based on the number of correct and incorrect classifications.

## Results

Training and testing of the algorithm was based on 18,000 hours of manually verified EEG data from mouse model of TLE. However, the nature of scalp EEG recording results in noisy data as indicated by **Figure 5**. To compensate for the prevalence of noise, we included various types of noise clips within our training dataset and testing dataset.

**Figure 5:**
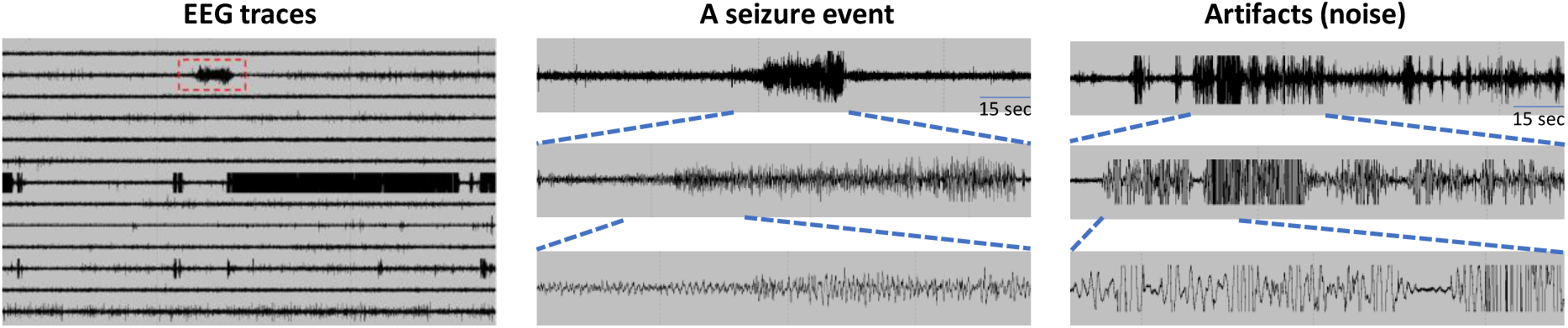
A representative sample of typical EEG events. A.) EEG traces recorded from 13 animals. The event highlighted in red represents a true, verified seizure. The rest are all noise/artifacts or normal brain activity. B.) Typical seizure event at three different scales. The top-level event represents the full seizure event, while subsequent EEG traces consist of zoomed-in fractions of the full seizure event. C.) Typical noise/artifact signal. The top-level event represents the full seizure event, and subsequent EEG traces represent more detailed slices of the noise signal.

To capture the global shape of the seizure and non-seizure events, each data clip was normalized and a simple average of each clip was taken, as seen by **Figure 2**. After transformation, differentiation between seizures and non-seizures could already be seen. Seizure events were marked by a gradual increase in electrical activity, while non-seizures events experienced sharp increases in electrical activity.

In cross-validation, our model trained off 64 clips consisting of 32 seizure clips and 32 non-seizure clips that were manually verified. The most optimal parameters for C and gamma were found to be 3.162 and 0.1, respectively, with a cross-validation score of 0.96 (**Figure 4**).

In the final assessment, 546 manually verified clips were tested throughout 5 trials (**Figure 6**). The outputted classification array was subtracted by the actual classifications to produce an array of 1s and 0s, where 1s represent incorrect detections and 0s represent correct detections. The average sensitivity to seizures was 40/42, or 95.24%. The average false positive rate was 6/504, or 1.2% giving us a specificity rate of 98.8%. The overall average accuracy of correct classification was 98.51% (**Figure 6**).

**Figure 6:**
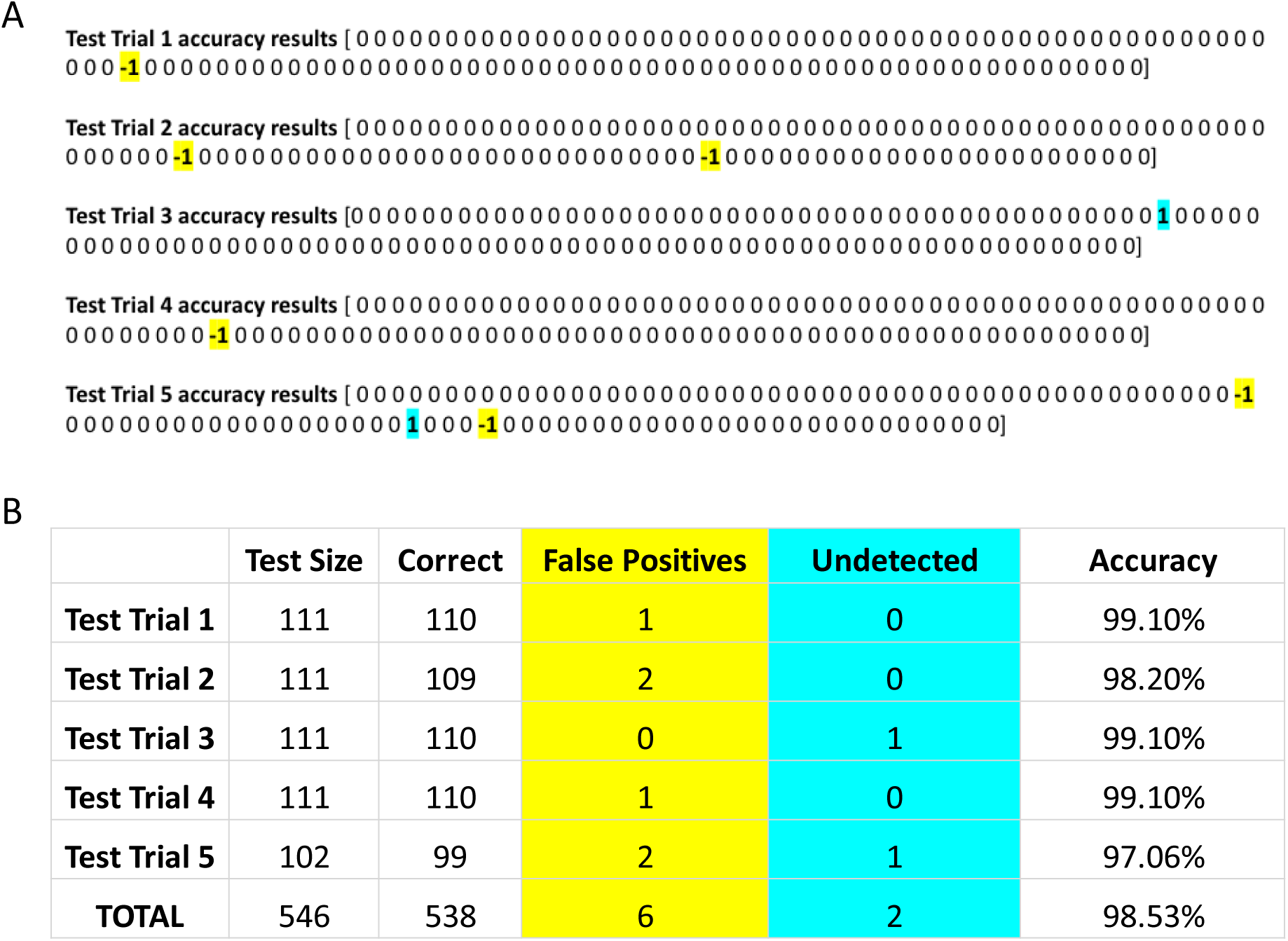
Testing results. Testing data set consisted of 42 seizures and 504 non-seizures (noise and other controls). Testing data was split into 5 groups. A.) Testing results for each trial. The outputted classification array of each data set was subtracted by the actual classifications to produce an array of 1s and 0s, where 1s represent incorrect detections and 0s represent correct detections. B.) Summary of testing results.

## Discussion

Our SVM classification was based on EEG data points that were processed and condensed to extract the global trends of each EEG signal. This process was designed to mimic human visual analysis of seizures, forgoing discrete values for larger, more notable trends in EEG clips. This novel method of seizure analysis achieved an accuracy rate of approximately 98.51%, comparable to that achieved by other algorithms (Saini and Dutta, 2017). While other implemented algorithms have used various high-level, computationally expensive methods of data transformation and feature extraction such as Fourier transformations and chaos theory (Baldassano et al., 2017; Saini and Dutta, 2017), the success of our novel model shows the potential of simple, broad trends of condensed EEG signals in classification. Specifically, our algorithm uses a small fraction of data points. This simple method could have an advantage over others in low-power platforms such as smartwatches, smartphones, and other wearables.

However, despite our novel methodology, our experiment still suffers from the same limitations as others: size of data that were tested. Our experiment, like other experiments (Baldassano et al., 2017; Liu et al., 2012; Saini and Dutta, 2017; Zhang and Chen, 2017), only used a fraction of the total amount of annotated data available. In addition, our experiment suffered from another common limitation to experimentation in the composition of our data. This experiment was tested using a data set with a seizure:noise ratio of 1:13. The real seizure:noise ratio is likely a thousand fold smaller, approximately 1:100 or even 1:10000. Thus, in theory, seizure detection rate around 98% is still not good enough for the real implementation of algorithms reported in the previous studies and presentation. A significantly higher accuracy is needed for the real implementation of an automated seizure detection system. As our approach is a significantly different from others, the combination of the methods proposed in this experiment as well as the those implemented by other researchers using other seizure features (i.e seizure spike-frequency, post-ictal depression etc.) could work synergistically to increase overall accuracy, making the system robust enough to handle a larger, more realistic data set.

Many laboratories have focused on developing novel algorithms for prediction of seizures. However, these algorithms often suffer significantly in low accuracy, and sacrifice computational cost, making these algorithms inapplicable in wearable devices. Our algorithm, on the other hand, is significantly more accurate and computationally more efficient than other seizure prediction algorithms through our data transformation strategy. The features that our algorithm analyzes are more robust than those of other algorithms, because while other algorithms focus on minute features within the data that fluctuate easily, our algorithm focuses on broad trends within the global shape of the data, allowing for a greater margin of error within our algorithm. In addition, while our algorithm does not predict seizures before they occur, our algorithm still has a powerful implication in epilepsy management, specifically in wearables. Our algorithm can be used on wearables for alerting emergency care (**Figure 7A**). Our algorithm has a particular niche in managing status epilepticus, one of the most dangerous types of seizure events that lasts between 10 – 30 minutes. Because status epilepticus is marked by multiple tonic-chronic seizures occurring in succession, our algorithm can potentially detect and deliver therapy within 1 min of status epilepticus (**Figure 7B**).

**Figure 7:**
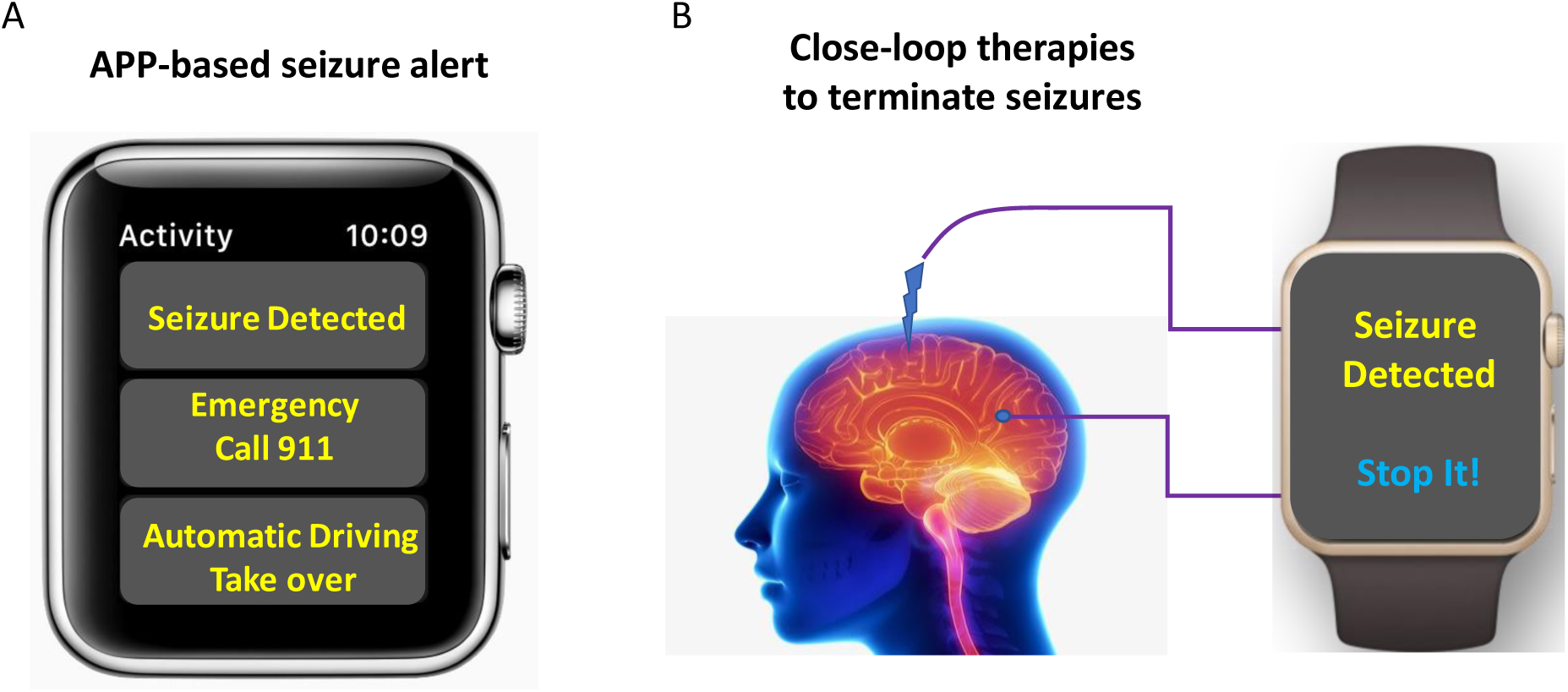
Diagram showing potential implications of our novel algorithm in seizure alert and on-site treatment for the severe forms of epilepsy.

## Conclusion

This novel model of seizure analysis focuses on broader trends, using less than 1/10 of data points. Yet, it achieved an accuracy rate of approximately 98.51%, comparable to that of other algorithms that use more computationally expensive methods of data analysis and require much more data. This theoretical model could be implemented in epilepsy research, allowing researchers to quickly comb through vast amounts of EEG data to find seizures. In addition, our computationally light-weight model for condensing EEG traces for global-trend analysis makes it more possible for implementation of automated seizure detection software on small, low RAM mobile devices such as wearables. Notably, the FDA recently approved a seizure detection app called embrace2 by Empatica. However, their algorithm detects seizures indirectly by monitoring heart rate and muscle contractions, whereas our novel algorithm detects seizures directly through analysis of brain activity. Thus, our algorithm may be more suitable for future wearable epilepsy management.

